# Sexually Dimorphic Circadian Regulation of Eclosion and Clock Gene Expression in a Migratory Moth *Spodoptera frugiperda*

**DOI:** 10.1101/2025.04.21.649798

**Authors:** Changning Lv, Yibo Ren, Viacheslav V. Krylov, Yumeng Wang, Yuanyuan Li, Weidong Pan, Gao Hu, Fajun Chen, Guijun Wan

## Abstract

The circadian clock orchestrates essential behavioral and molecular processes, including the timing of eclosion, one of the most tractable and ecologically relevant outputs of the circadian system. Understanding eclosion timing may offer insights into the circadian clock mechanisms that underlie migratory timing. Here, we characterize the diel and circadian patterns of eclosion and core clock gene expression in the fall armyworm (FAW), *Spodoptera frugiperda*, a globally distributed migratory moth. Using a custom-designed eclosion monitoring system under both 14 hours light-10 hours dark (L14: D10) and constant darkness (DD) conditions, we observed clear diel eclosion rhythms, peaking shortly after lights-off under L14: D10. Under DD, these rhythms became delayed and damped over three consecutive days, consistent with circadian control. Males exhibited more dispersed emergence patterns and distinct eclosion distributions than females under L14: D10 and DD, suggesting sexually dimorphic timing. Gene expression profiling revealed rhythmic oscillations of five canonical clock genes, *cyc*, *clk*, *tim*, *per*, *cry2*, with sex-dimorphic differences in mesor, amplitude, or phase, particularly the mesors of males are higher than those of females in five clock genes under L14: D10.

These results provide strong evidence for sexually dimorphic circadian regulation at both behavioral and molecular levels in a migratory insect that recently invaded China, suggesting that sex-dimorphic circadian architecture may contribute to allochronic speciation and ecological strategies such as eclosion and migratory timing in a novel environment. Leveraging the established circadian eclosion rhythm assay, which captures core features of clockwork mechanisms, further investigation may reveal how these sex-based differences contribute to the evolution of circadian function and adaptive traits.

## Introduction

Circadian timing enables organisms to synchronize their physiology and behavior with the daily cycle, allowing them to anticipate and adapt to environmental fluctuations (Hardin, 2011; Reppert and Weaver, 2002). One of the earliest circadian rhythms to be studied is the rhythm of adult emergence (eclosion) in holometabolous insects (Bunning, 1935; Kalmus, 1935; Saunders, 2002). Eclosion marks the completion of metamorphosis and is typically gated to dawn or dusk, depending on the species. This event exhibits circadian rhythmicity at the population level, persisting under constant conditions and demonstrating temperature compensation—hallmarks of endogenous circadian control (Numata and Tomioka, 2023; Pittendrigh, 1954; Pittendrigh and Skopik, 1970). Under natural conditions, synchrony between the internal circadian system and external environmental cues such as photoperiod, the principal zeitgeber or temporal cue, facilitates the optimal timing of eclosion in a species-specific manner, suggesting that the circadian system restricts emergence to the most ecologically advantageous periods of the day (Merlin, 2009).

Recent work in *Drosophila* has shown that the circadian clock gates eclosion by rhythmically modulating the terminal steps of metamorphosis (Mark et al., 2021; Wegener et al., 2024). The circadian clock is governed by a conserved transcriptional–translational feedback loop involving core clock genes shared across most insects, including *cycle* (*cyc*), *clock* (*clk*), *timeless* (*tim*), *period* (*per*), and *cryptochrome* (*cry*) (Pilorz et al., 2018; Tomioka and Matsumoto, 2015). Oscillations in the expression of these genes within central and peripheral tissues regulate downstream clock-controlled genes, which in turn orchestrate overt rhythmic behaviors like eclosion. Studies have demonstrated that disruption of these core clock genes alters the timing and rhythmicity of eclosion (Dushay et al., 1990; Konopka and Benzer, 1971; Sehgal et al., 1994; Zhang et al., 2017), establishing eclosion as one of the most accessible and ecologically relevant behavioral outputs of the circadian clock. Utilizing eclosion as a model behavior provides a powerful framework for dissecting circadian clock mechanisms and understanding their contributions to insect adaptation (Iiams et al., 2019, 2024; Miyazaki et al., 2016; Myers, 2003; Qiu et al., 2023).

Circadian clocks play a critical role in regulating the physiology and behavior of migratory insects, such as mediating the seasonal induction of migratory traits, regulating flight rhythms, determining reproductive state, and facilitating orientation during migration (Ji et al., 2022; Merlin et al., 2020; Merlin and Liedvogel, 2019). Besides, precise circadian timing becomes especially important for migrants as individuals must adjust rapidly to shifting photoperiods and environmental cues across broad geographic gradients. Thus, the circadian system may serve as a central temporal framework that integrates endogenous rhythms with environmental signals, enabling migratory insects to time their departure, orientation, and arrival in synchrony with ecological opportunities, although the underlying mechanisms remain to be further elucidated (Merlin and Liedvogel, 2019).

One such migratory species is the fall armyworm, *Spodoptera frugiperda*, a globally distributed noctuid moth and major agricultural pest. Originating from the tropical and subtropical areas of the Americas, this pest was first detected in the African regions of Nigeria and Ghana in 2016 (Goergen et al., 2016). Exploiting its formidable migratory capacity, it rapidly expanded its invasive range across Sub-Saharan Africa, the Middle East, much of Asia, and Oceania (Kenis et al., 2022; Tay et al., 2023). By December 2018, it had reached Jiangcheng County, Yunnan Province, China (Hui et al., 2020; Li et al., 2019; Sun et al., 2021). From year-round breeding zones in South China and Southeast Asia, *S. frugiperda* spreads northward across East Asia during the spring and summer months (Huang et al., 2022; Jiang et al., 2022; Wu et al., 2021; Zhang et al., 2023). Our recent flight simulation studies revealed flight orientations directed north-northwest in spring and southwest in autumn, which would promote seasonal forward and return migrations in East Asia. Photoperiod has been identified as the principal environmental cue driving this seasonal shift (Chen et al., 2023), suggesting a potential role for the circadian clock in the temporal regulation of migratory orientation (Denlinger et al., 2017; Jin et al., 2023). Another well-known circadian-regulated phenotype in *S. frugiperda* could be the allochronic mating behavior observed between two morphologically identical but genetically distinct strains, which is a trait unique to populations in the United States (Hanniger et al., 2017; Miller et al., 2024). A recent simulation study suggested that sex-dimorphic expression of circadian rhythms can facilitate allochronic speciation (van Doorn et al., 2024), implying the most likely presence of sexually dimorphic circadian clocks in native populations within the United States. In contrast, invasive *S. frugiperda* populations exhibit a largely homogeneous genetic background distinct from their native counterparts, likely due to hybridization between the strains during the invasion process (Wang et al., 2024). Regardless of genetic background, experimental evidence for sex-dimorphic circadian rhythms beyond mating behavior in *S. frugiperda* remains limited, leaving a critical gap in our understanding of how circadian mechanisms may shape strain divergence and ecological adaptation.

Using *S. frugiperda* as a model, this study aims to investigate the circadian regulation of adult emergence (eclosion), with a particular focus on potential sexual dimorphism in behavioral rhythms and clock gene expression. We characterized diel and circadian patterns of eclosion in males and females under both light-dark (LD) and constant darkness (DD) conditions using a customized monitoring system. To complement behavioral observations, we also quantified temporal expression profiles of core clock genes (*cyc*, *clk*, *tim*, *per*, *cry2*) in adult heads across sexes and lighting regimes. By integrating phenotypic and molecular data, we explored whether sex-dimorphic circadian architectures underlie differences in eclosion timing and may reflect broader implications for strain divergence and migratory timing. These findings will advance our understanding of migratory insect chronobiology and the evolutionary plasticity of the circadian clock, while also informing predictive frameworks for the management of migratory insect pests.

## Material and methods

### Insect stock

The fall armyworm *S. frugiperda* strain was originally collected from a maize field in Yuanjiang County, Yuxi City, Yunnan Province, China (23.604°N, 101.977°E) during their migration season. They were housed in the laboratory for multiple generations under a summer-like photoperiod of 14 hours light: 10 hours dark (L14: D10) at 26 ± 1 ℃ and 70 ± 5 % humidity to mimic a summer-like condition when they cause the most crop damage and conduct partial migration outdoors from south to north (Li et al., 2019; Wu et al., 2021). The 1st to 3rd instar larvae of *S. frugiperda* were reared with fresh sweet maize leaves (Huajintian No. 2 sweet maize) in plastic containers measuring 40 × 20 × 15 cm. The 4th to 6th instar larvae were placed individually in *Drosophila* vials (2.4 cm bottom diameter, 9.5 cm height) with sufficient artificial diet (Li et al., 2019). The vials were sterilized, and new artificial diets were added every three days to maintain a clean environment and ensure adequate food supply until pupation. The pupation date was recorded, and the sex was identified before the pupae were transferred to new numbered *Drosophila* vials containing a moisturized cotton ball to maintain the relative humidity. Adults of *S. frugiperda* were fed using a 10% honey solution daily.

### Diel eclosion behavior assay under the summer-like LD cycle

Eclosion behavior was performed with a customized apparatus for insect eclosion behavior monitoring (Fig. 1A). *S. frugiperda* larvae were raised in summer-like LD (i.e., L14: D10) in an incubator (HPG280H, Ningbo Jiangnan Ltd., Ningbo, China) at 26 ± 1℃ and 70 ± 5 % humidity through pupation. Zeitgeber time 0 (ZT0) was defined as the time of light onset, and ZT14 as the time of dark onset. The pupal duration under summer-like and constant darkness (i.e., Dark-Dark, DD) conditions ranges from 10.04±0.10 days to 11.42±0.10 days. For acclimation purposes, 7-day-old pupae were transferred gently to a numbered glass tube (2.0 cm outside diameter, 7.0 cm height, 0.1 cm thickness) with open ends during the dark period 1 hour before the beginning of the next LD or DD cycle, using a moisturized cotton ball to seal both ends. All tubes were placed on the apparatus for diel adult eclosion monitoring by an infrared-based camera system under LD conditions (Fig. 1). Remove the newly emerged adults from the apparatus daily under red light at 8:30 am (ZT0). Diel eclosion rhythms were analyzed using the eclosion profile over five consecutive LD cycles.

**Figure 1.**
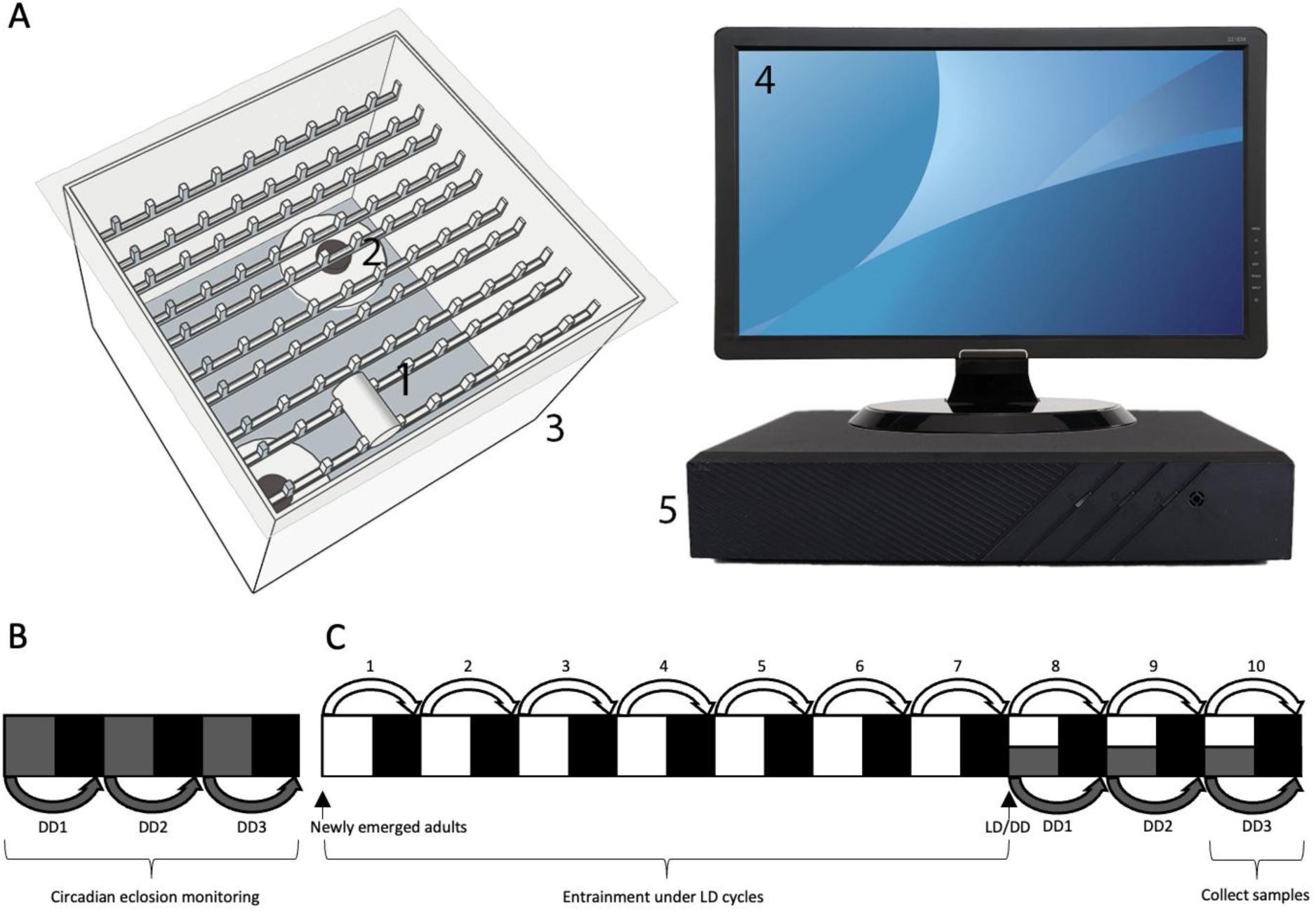
The assay for eclosion rhythm monitoring and adult tissue collection of *Spodoptera frugiperda*. (A) The real-time eclosion monitoring apparatus for insects. Key components include: 1. Glass tube; 2. Camera; 3. Acrylic stand; 4. Screen; 5. Video recorder. Cameras are connected to a video recorder through network cables. Granted Chinese patent: ZL202120866706.1. (B) Circadian eclosion monitoring assay under the 1st (DD1), 2nd (DD2), and 3rd (DD3) dark-dark (DD) cycles. Dark grey and black bars represent subjective day and night, respectively. (C) Entraining adults under 14 hours light: 10 hours dark (LD) for seven cycles, followed by sample collection for gene expression analysis after two additional LD or DD cycles. Heads of *S. frugiperda* were collected every 4 hours under LD conditions and every 3 hours under DD3 conditions over 24 hours.

### Circadian eclosion behavior assay under constant darkness

Pupae were transferred to DD from LD condition 1 (DD1), 2 (DD2), and 3 (DD3) days before the time of adult eclosion for circadian eclosion monitoring (Fig. 1B). Circadian eclosion behavior was performed as described in LD condition but under constant darkness. Circadian time 0 (CT0) is defined as the beginning of the subjective day, and CT14 is the beginning of the subjective night. Remove the newly emerged adults from the apparatus at 8:30 am (CT0) under red light daily. Circadian eclosion rhythms were analyzed using the eclosion profile over three consecutive DD cycles.

### Tissue collection

For RNA extractions of the LD condition, heads from newly emerged female or male adults entrained to ten LD cycles were collected every 4 hours over 24 hours, with three heads pooled per sample. For RNA extractions of DD conditions, heads from female or male adults entrained to seven LD cycles were collected every 3 hours over 24 hours under red light on the third day (DD3; 10-day-old) of transfer into DD, with three heads pooled per sample. The samples were stored at-80°C until further use.

### RNA isolation and cDNA synthesis

Three biologically independent head pools were used for each group based on sex and sampling time. Total RNA was extracted from these pooled samples using TRIzol® (Invitrogen; Thermo Fisher Scientific, Inc., Waltham, MA, USA). The extracted RNA samples were individually analyzed for quality and quantity using a NanoDrop2000 (Thermo Fisher Scientific, Inc., Waltham, MA, USA). Before reverse transcription, the integrity of each total RNA sample was assessed by electrophoresis in a 1% agarose gel. Subsequently, cDNA was synthesized from 100 ng of total RNA in a 20 μl reaction, utilizing the PrimeScript RT reagent kit supplemented with a gDNA Eraser (Takara Bio Inc., Dalian, China).

### Gene expression analysis

Five target genes, *cyc, clk, tim, per, and cry2,* were selected for gene expression analysis using a quantitative real-time polymerase chain reaction (qRT-PCR) assay. Two reference genes, *EF1α* and *rpl32*, were evaluated for stability with a standard deviation value lower than 1 (Rajarapu et al., 2011; Y. Zhang et al., 2022) across sexes to ensure normalization reliability (Fig. S1). Primers specific to each gene were designed individually using the Oligo 7 software (Molecular Biology Insights, Inc., Cascade, CO, United States). The synthesis of primers (Table 1) was completed by GeneScript Biotechnology Co., Ltd. (China) and tested as previously described (Y. Zhang et al., 2023). The qRT-PCR was conducted on an Applied Biosystems® 7500 Fast Real-Time PCR System (Thermo Fisher Scientific, Inc., Waltham, MA, USA) using SYBR Premix Ex Taq (Tli RNaseH Plus; Takara Bio Inc., Dalian, China). The reactions were conducted in a final volume of 20 μl (including 2 μl of a 1/10 dilution of the cDNA template and primers in a final concentration of 200 nM) with the following conditions: an initial 30 s step of 95°C followed by 40 denaturation cycles at 95°C for 5 s and primer annealing at 60°C for 34 s. The *EF1α* and *rpl32* were used as the reference genes (Zhang et al., 2022), and the 2^−ΔΔCt^ method (Ct, cycle threshold) was applied to evaluate the relative expression levels (Livak and Schmittgen, 2001). Three biological replicates were used for statistical comparison between groups.

**Table 1.**
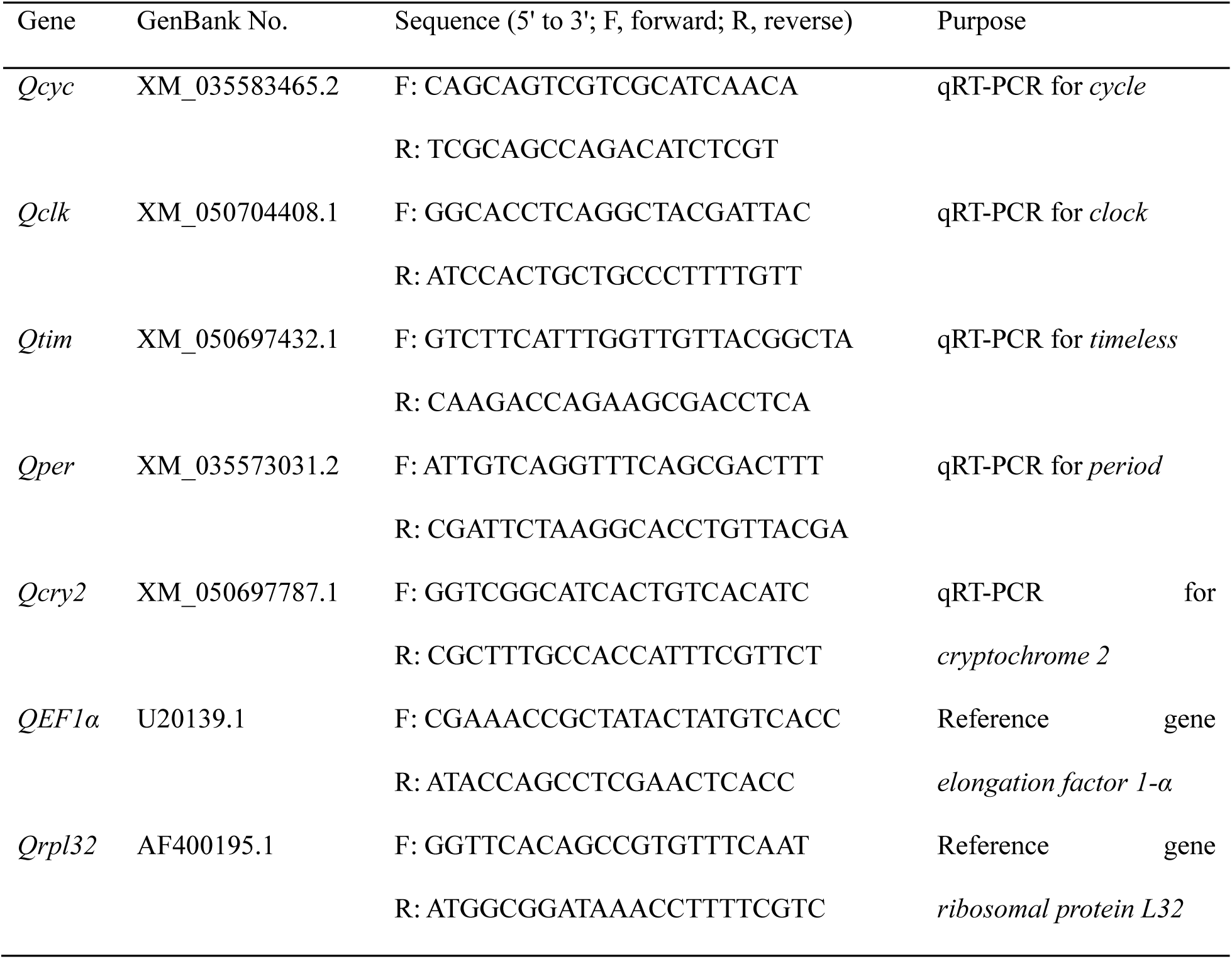
Primers of the five target genes and two reference genes.

### Statistical analysis

The Kolmogorov-Smirnov test (N>50) and Shapiro-Wilk test (N≤50) were used to test for normality (*P* > 0.05), and Levene’s test for the homogeneity of variances (*P* > 0.05), before an analysis of variance (ANOVA). The non-parametric Mann-Whitney test and exact Kolmogorov-Smirnov (KS) test maximum difference were used to test the difference in eclosion distribution. One-way ANOVA was used to test the gene expression difference between sexes at a given ZT or circadian time CT. Multivariate tests were applied to test gene expression differences between time points for females and males. Two-way ANOVA was used to test the interaction effects between time points and sex under the same photoperiod condition. The above tests were conducted with SPSS 26. Rhythm characteristics, including mesor (midline-estimating statistic of rhythm), amplitude, and phase between different photoperiod condition-sex combinations, were determined using the CircaCompare algorithm (Parsons et al., 2019). Data were plotted using GraphPad Prism 8.

## Results

### Diel and circadian eclosion rhythms of S. frugiperda entrained to summer-like photoperiod

The average eclosion peak under LD conditions (pooled data) occurred at the 1st hour after lights-off (ZT14-ZT15), with 58% of females and 40% of males emerging over 24 hours (Fig. 2A). The average eclosion peak under DD conditions (pooled data) occurred at the 2nd hour after the onset of subjective dark (CT15-CT16), with 27% of females and 19% of males emerging over 24 hours. Another average eclosion peak under DD conditions was also observed between CT17-CT18 in male adults (Fig. 2B). Under DD conditions, the average eclosion peak of *S. frugiperda* exhibited a phase delay of approximately one hour compared to those observed under LD conditions. The eclosion peak is sharper and more pronounced under LD conditions, whereas in DD, it is more gradual and phase-delayed (Fig. 2A, B). Moreover, across successive DD cycles (DD1 to DD3), the eclosion peak progressively shifts later each day (Fig. 2C). Females took fewer circadian cycles to complete eclosion, with only one sample emerging in DD3 for the same generation compared to males. Therefore, we excluded the eclosion data for females in DD3 from the subsequent statistical comparison between groups.

**Fig. 2.**
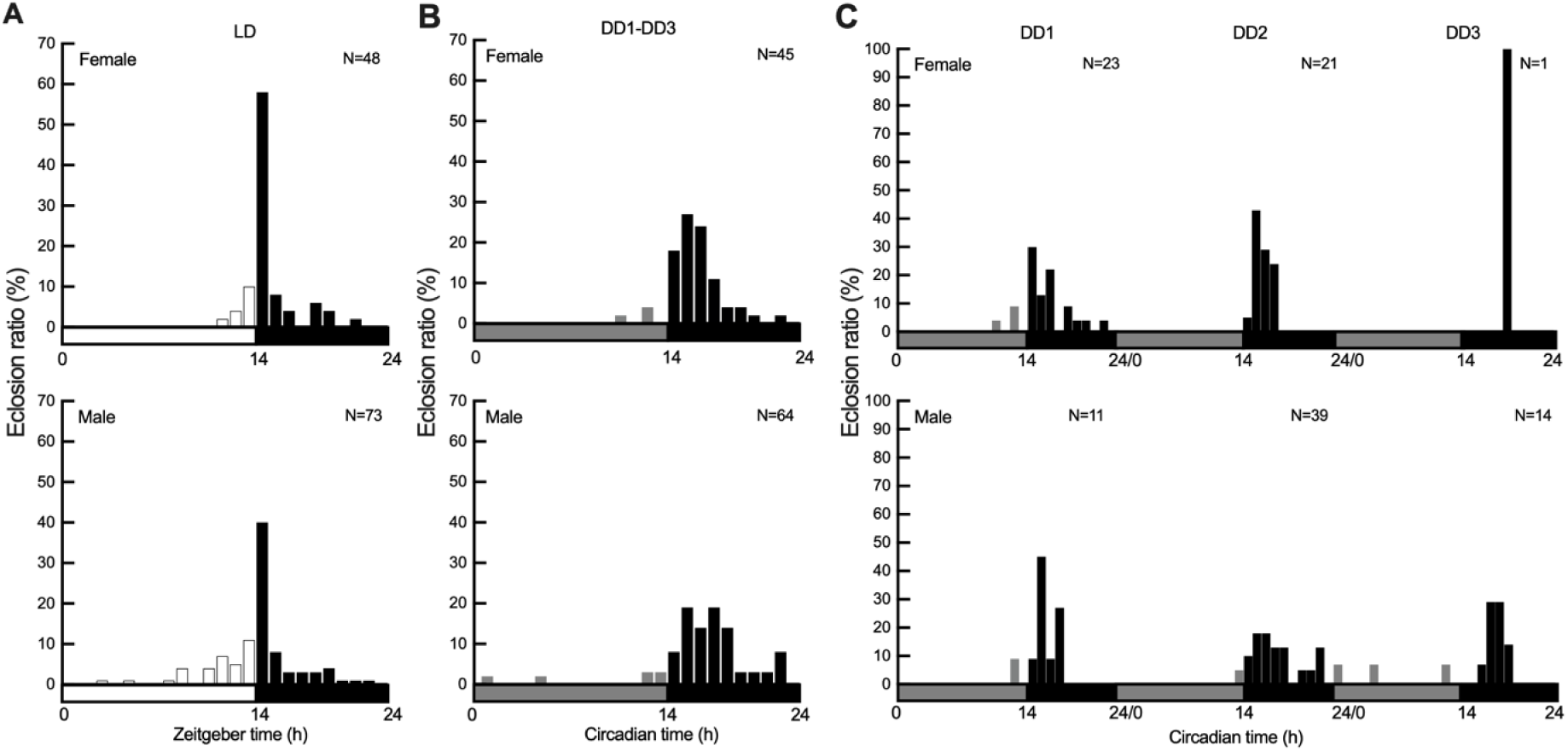
Diel and circadian eclosion rhythms of *S. frugiperda* entrained to summer-like photoperiod. (A) Pooled diel eclosion rhythm over consecutive five days of recording under 14-hour light: 10-hour dark (LD) conditions. (B) Pooled circadian eclosion rhythm of females and males over consecutive three days of recording under constant darkness (DD) after entrainment to LD. (C) Profiles of female and male adult eclosion across the 1st, 2nd, and 3rd days of constant darkness (DD1, DD2, and DD3, respectively) after entrainment to LD. Data are binned in 1-h intervals. Horizontal bars show objective day (white) and night (black) in panel (A) and subjective day (gray) and night (black) in Panel (B) and (C). F, Female; M, Male. The overall distribution of eclosion times for males and females was compared using the exact Kolmogorov-Smirnov test maximum difference: Pooled LD-F versus Pooled LD-M, *P* = 0.059; Pooled DD-F versus Pooled DD-M, *P =* 0.019; Pooled LD-F versus Pooled DD-F, *P <* 0.001; Pooled LD-M versus Pooled DD-M, *P <* 0.001; DD1-F versus DD1-M, *P* = 0.362; DD2-F versus DD2-M, *P* = 0.017; DD1-F versus DD2-F, *P =* 0.010; DD1-M versus DD2-M, *P =* 0.085; DD1-M versus DD3-M, *P =* 0.058; DD2-M versus DD3-M, *P =* 0.334. *P* < 0.05 suggests significant effects on the overall distribution pattern of eclosion times, including characteristics such as variability and distribution shape, beyond just the median.

### Distribution of adult eclosion between sexes and photoperiod conditions

To gain a more detailed understanding of the overall distributions of adult eclosion in *S. frugiperda*, the exact Kolmogorov-Smirnov (KS) test was applied to examine differences in eclosion distributions between photoperiod conditions and sexes. A highly significant difference was found when comparing eclosion distributions between pooled LD and DD conditions for both females (*P* < 0.001) and males (*P* < 0.001), indicating a marked shift in timing and pattern under constant darkness, consistent with a circadian-driven free-running period. Additionally, marginal and significant differences were detected between female and male eclosion rhythms under pooled LD (*P* = 0.059) and DD conditions (*P* = 0.019), respectively, suggesting sexually dimorphic circadian regulation of eclosion timing (Fig. 2A, B). Due to the limited sample size of females on DD3, statistical comparisons for this group were not performed. Further analysis of consecutive D1-D2 for females and D1-D3 for males revealed significant sex-dimorphic differences in eclosion distribution on DD2 (*P* = 0.017), consistent with the pooled DD condition results. In contrast, no significant difference was observed for DD1 (*P* = 0.362), likely due to a small sample size. Within-sex comparisons across different DD days revealed marginal to significant effects, with males showing marginal differences between DD1 and DD2 (*P* = 0.085) and DD1 and DD3 (*P* = 0.058), while females exhibited a significant effect between DD1 and DD2 (*P* = 0.010). No significant difference was found for males between DD2 and DD3 (*P* = 0.334).

Given that the Mann-Whitney (MW) test is sensitive to differences in central tendency (median), while the KS test captures overall distributional differences more effectively, the MW test was further employed to specifically assess differences in the central tendency of adult *S. frugiperda* eclosion per hour relative to ZT0/CT0 between sexes and photoperiod conditions. A marginal difference was observed between sexes under LD conditions (*P* = 0.095), whereas significant differences were detected between sexes under DD conditions (*P =* 0.032). Additionally, highly significant differences were observed when comparing eclosion distributions between pooled LD and pooled DD conditions for both females and males (*P* < 0.001; Fig. 3). The results of the MW test were largely consistent with those of the KS test, suggesting that shifts in central tendency contribute to the overall distribution differences. The marginal difference detected by the MW as well as KS test under LD conditions may reflect a slight trend in median eclosion times, likely influenced by the higher frequency of male eclosion during the daytime (Fig. 2A; Fig. 3). We further examined circadian eclosion dynamics over three consecutive DD days. Consistent with the KS test, the MW test revealed no significant differences between sexes for DD1 (*P* = 0.744) but a marginal difference for DD2 (*P* = 0.087). For males, no significant difference was found between DD2 and DD3 (*P* = 0.943), while marginal differences were observed between DD1 and DD2 (*P* = 0.061) and between DD1 and DD3 (*P* = 0.058). However, in contrast to the KS test, the MW test detected no significant differences between DD1 and DD2 for females (*P* = 0.216), suggesting that the significant difference observed in the KS test was primarily due to differences in the distribution shape and spread, rather than in location (Fig. 3).

**Fig. 3.**
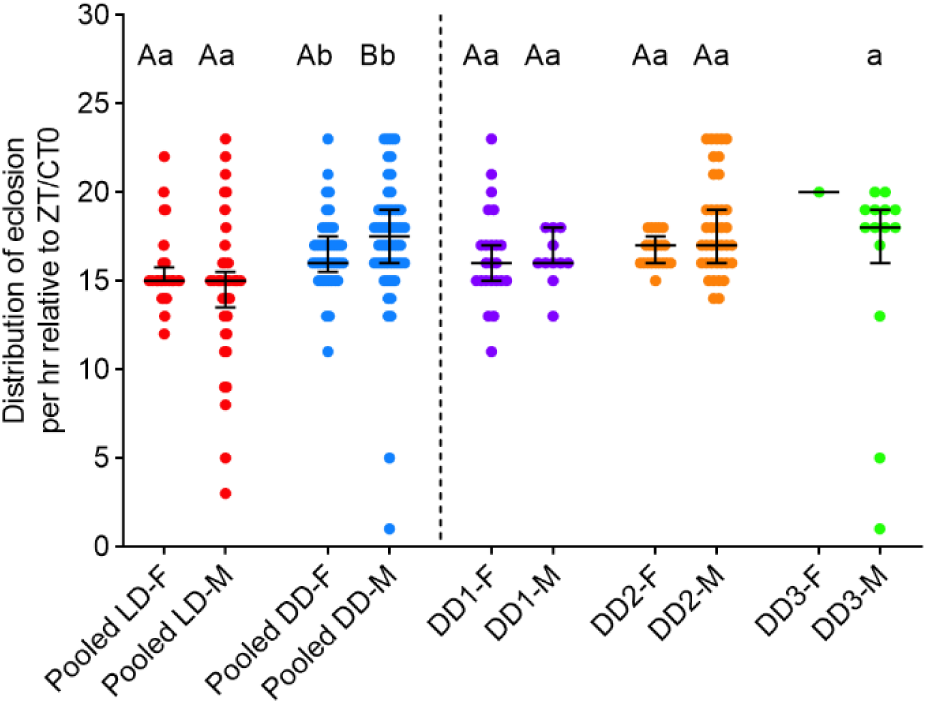
Central tendency comparison for sex differences in eclosion rhythm of *S. frugiperda* under 14-hours light: 10-hours dark (LD) and constant darkness (DD) conditions. Red, blue, purple, orange, and green dots represent data from five consecutive days under 14-hour light: 10-hour dark (Pooled LD), three consecutive days under constant darkness (Pooled DD), the 1st (DD1), 2nd (DD2) and 3rd (DD3) day of constant darkness, respectively. F, Female; M, Male. Mann-Whitney test: Pooled LD-F versus Pooled LD-M, *P* = 0.095; Pooled DD-F versus Pooled DD-M, *P =* 0.032; Pooled LD-F versus Pooled DD-F, *P <* 0.001; Pooled LD-M versus Pooled DD-M, *P <* 0.001; DD1-F versus DD1-M, *P* = 0.744; DD2-F versus DD2-M, *P* = 0.087; DD1-F versus DD2-F, *P =* 0.216; DD1-M versus DD2-M, *P =* 0.061; DD1-M versus DD3-M, *P =* 0.058; DD2-M versus DD3-M, *P =* 0.943. Uppercase and lowercase letters indicated significant differences between sexes for the same photoperiod condition (LD or DD), and between the photoperiod conditions for females or males by Mann-Whitney test at *P <* 0.05, respectively. Solid lines represent the median value, and error bars represent the interquartile range. *P* < 0.05 indicates a significant difference in the central tendency (median) of eclosion times between treatments.

### Transcript expression of circadian genes in the heads of S. frugiperda under LD and DD conditions

To investigate the expression of core circadian genes between sexes and photoperiod conditions, we conducted qRT-PCR to examine the levels of *cyc*, *clk*, *tim*, *per*, and *cry2* in the heads of *S. frugiperda* collected at 3-or 4-hour intervals. After 7 days of entrainment to an LD cycle, 10-day-old adults were sampled on the 3rd day of DD to ensure a clearer observation of the free-running rhythm of the molecular circadian clock.

Under the LD condition, significant interactions between sex and time were observed for *cyc* (*P =* 0.013; Fig.4A) *and marginal difference for clk* (*P =* 0.079; Fig.4B) by two-way ANOVA. One-way ANOVA between sexes for the same time point revealed significant sex differences in gene expression levels at: ZT5 (*P* = 0.002), ZT9 (*P* = 0.008), and ZT17 (*P* = 0.007) for *cyc* (Fig.4A); ZT5 (*P* = 0.034), ZT9 (*P* = 0.012), and ZT17 (*P* < 0.001) for *clk* (Fig.4B); ZT1 (*P* = 0.025) and ZT5 (*P* = 0.021) for *tim* (Fig.4C); ZT5 (*P* = 0.042) and ZT17 (*P* = 0.048) for *per* (Fig.4D); and ZT9 (*P* = 0.029) and ZT17 (*P* = 0.001) for *cry2* (Fig.4E).

**Fig. 4.**
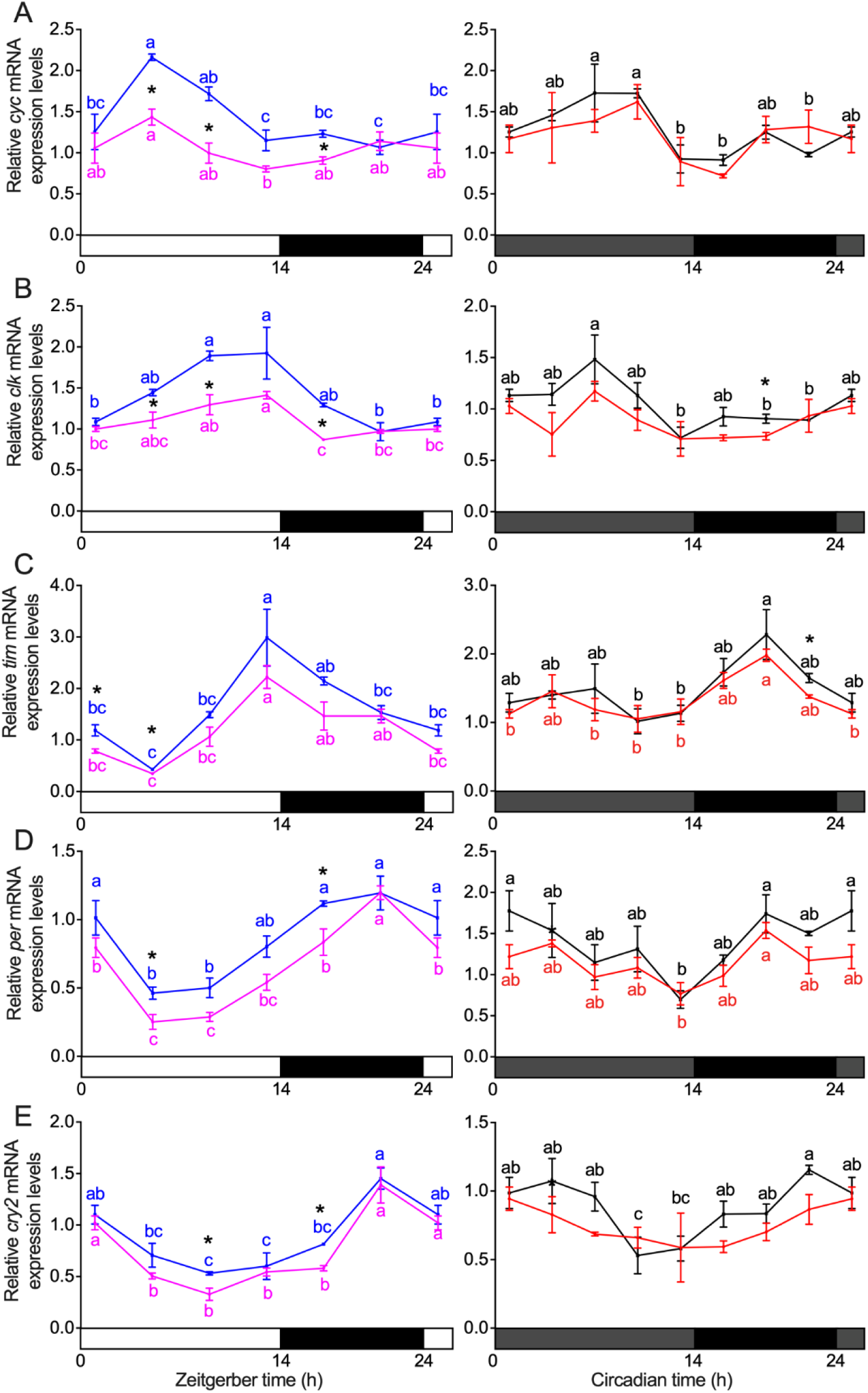
The expression of circadian genes of *cyc* (A), *clk* (B), *tim* (C), *per* (D), and *cry2* (E) in the heads of adults under 14 hours light: 10 hours dark (LD) and the 3rd day of constant darkness (DD) after 7 days of adult entrainment to LD. Horizontal bars show objective day (white) and night (black) in the left panels and subjective day (gray) and night (black) in the right panels. Pink and blue lines indicate females (LD-F) and males (LD-M) sampled under LD conditions, and red and black lines indicate females (DD-F) and males (DD-M) sampled under the 3rd day of DD after 7 days of adult entrainment to LD. Two-way ANOVA, interaction sex × time: (A) *cyc*: LD, *P =* 0.013; DD, *P* = 0.819. (B) *clk*: LD, P = 0.079; DD, *P* = 0.783. (C) *tim*: LD, *P* = 0.45; DD, *P* = 0.928. (D) *per*: LD, *P* = 0.51; DD, *P* = 0.83. (E) *cry2*: LD, *P* = 0.818; DD, *P* = 0.503. One-way ANOVA between sexes for the same time point (only significant differences were reported): (A) *cyc*: ZT5, *P* = 0.002; ZT9, *P* = 0.008; ZT17, *P* = 0.007. (B) *clk*: ZT5, P = 0.034; ZT9, *P* = 0.012; ZT17, *P* < 0.001; CT19, *P* = 0.045. (C) *tim*: ZT1, P=0.025; ZT5, *P* = 0.021; CT22, *P* = 0.019. (D) *per*: ZT5, *P* = 0.042; ZT17, *P* = 0.048. (E) *cry2*: ZT9, P = 0.029; ZT17, *P* = 0.001. The interactions between sex and time for each photoperiod condition were tested by two-way ANOVA at *P* < 0.05. The differences between sexes for the same time point were tested by one-way ANOVA at *P* < 0.05. Different lowercase letters indicated significant differences among different time points for the same sex under the same photoperiod condition by post hoc Tukey HSD test at *P* < 0.05. * *P* < 0.05. Solid lines represent the mean value, and error bars represent the standard error of the mean (SEM).

Under the DD condition, no significant interactions between sex and time were found for all tested circadian genes (Fig.4). Significant sex differences in gene expression levels were observed at CT19 (*P* = 0.045) for *clk* (Fig.4B); CT22 (*P* = 0.019) for *tim* (Fig.4C).

The presence of rhythmicity was found to be significant for most selected core circadian genes in both female and male adults under different photoperiod conditions, except female *cyc* and *cry2* under DD condition as determined by CircaCompare and confirmed by post hoc Tukey HSD test at *P* < 0.05 (Fig.4; Table 2). Because CircaCompare requires rhythmicity in both groups for valid comparison, parameter estimation was not possible when one sex lacked oscillation. This accounts for the omission of rhythm-related indices for cyc and cry2 under DD conditions in the sex-based comparisons in Table 3.

We further analyzed the mesor (midline-estimating statistic of rhythm), amplitude, and phase differences of selected core circadian genes between sexes under LD and DD conditions using CircaCompare. Significant and marginal differences in mesor were observed between females and males for *cyc* (-0.38, P < 0.001), *clk* (-0.32, P < 0.001), *tim* (-0.40, P = 0.014), *per* (-0.20, P < 0.001) and cry2 (-0.14, P = 0.048) under LD conditions. Under DD conditions, significant mesor differences were found for *clk* (-0.17, P = 0.012) and *per* (-0.22, P = 0.024). Moreover, significant differences in amplitude were observed between females and males for *clk* (-0.29, *P* = 0.0038) under LD conditions. Notably, for phase differences between females (estimated peak at ZT2.96) and males (estimated peak at ZT6.33; Table 3), we found significant differences for *cyc* (-3.37, *P* = 0.044; Table 3) under LD conditions.

**Table 2.** Presence of rhythmicity (*p*-value) of five circadian genes for different genders and light conditions using the CircaCompare method.

|  |  | *cyc* | *clk* | *tim* | *per* | *cry2* |
| --- | --- | --- | --- | --- | --- | --- |
| Female | LD | 0.0071** | <0.001*** | <0.001*** | <0.001*** | <0.001*** |
|  | DD | na | 0.044* | 0.0058** | 0.010* | na |
| Male | LD | <0.001*** | <0.001*** | <0.001*** | <0.001*** | <0.001*** |
|  | DD | <0.001*** | 0.0010** | 0.0059** | 0.0021** | <0.001*** |

**Note:** LD, 14-hours light: 10-hours dark; DD: constant darkness (DD) after 7 days of adult entrainment to LD.

**Table 3.** Differences in rhythmic parameters across groups using the CircaCompare method.

| Gene | Comparison | Mesor difference | Amplitude difference | Phase difference |
| --- | --- | --- | --- | --- |
| | Groups | ($P$ value) | ($P$ value) | ($P$ value) |
| *cyc* | LD-F vs. LD-M | -0.38 (<0.001<sup>***</sup>) | -0.24 (0.057) | -3.37 (0.044<sup>*</sup>) |
|  | LD-F vs. DD-F | na | na | na |
|  | DD-F vs. DD-M | na | na | na |
|  | LD-M vs. DD-M | na | na | -0.16 (0.90) |
| *clk* | LD-F vs. LD-M | -0.32 (<0.001<sup>***</sup>) | -0.29 (0.0038<sup>**</sup>) | -0.64 (0.60) |
|  | LD-F vs. DD-F | na | na | 5.59 (0.0069<sup>**</sup>) |
|  | DD-F vs. DD-M | -0.17 (0.012<sup>*</sup>) | -0.10 (0.27) | -1.12 (0.58) |
|  | LD-M vs. DD-M | na | na | 5.11 (<0.001<sup>***</sup>) |
| *tim* | LD-F vs. LD-M | -0.40 (0.014<sup>*</sup>) | -0.27 (0.22) | 0.42 (0.68) |
|  | LD-F vs. DD-F | na | na | -3.86 (0.015*) |
|  | DD-F vs. DD-M | -0.13 (0.24) | -0.094 (0.55) | -0.65 (0.72) |
|  | LD-M vs. DD-M | na | na | -4.93 (0.0021**) |
| *per* | LD-F vs. LD-M | -0.20 (<0.001***) | 0.055 (0.42) | 0.45 (0.46) |
|  | LD-F vs. DD-F | na | na | -3.21 (0.017*) |
|  | DD-F vs. DD-M | -0.22 (0.024*) | -0.17 (0.22) | -0.41 (0.83) |
|  | LD-M vs. DD-M | na | na | -4.07 (0.0038**) |
| *cry2* | LD-F vs. LD-M | -0.14 (0.048*) | 0.037 (0.70) | -0.050 (0.95) |
|  | LD-F vs. DD-F | na | na | na |
|  | DD-F vs. DD-M | na | na | na |
|  | LD-M vs. DD-M | na | na | -2.42 (0.021*) |

**Note:** Rhythmicity was found to be significant for all genes in both female and male adults under different photoperiod conditions by CircaCompare at $P < 0.05$ except female *cyc* and *cry2* under constant darkness (DD) condition. A negative sign indicates that the latter condition is reduced or delayed compared to the former. Data are presented with two significant digits after the decimal point. LD-F or LD-M, heads sampled from female or male adults under 14 hours light: 10 hours dark (LD); DD-F or DD-M: heads sampled from female or male adults under the 3rd day of DD after 7 days of adult entrainment to LD. \* $P < 0.05$ , \*\* $P < 0.01$ , \*\*\* $P < 0.001$ . For the comparison of group A vs. group B, group B is used as the reference group. Thus, the difference is calculated as A - B (e.g., Phase difference = $\phi_A - \phi_B$ ).

We also compared phase differences between LD and DD conditions and found significant differences in both females and males for *clk* (females: 5.59, *P* = 0.0069; males: 5.11, *P* < 0.001), *tim* (females:-3.86, *P* = 0.015; males:-4.93, *P* = 0.0021), *per* (females:-3.21, *P* = 0.017; males:-4.07, *P* = 0.0038), and *cry2* (males:-2.42, *P* = 0.021) (Table3; see Table 4 for peak time hours).

**Table 4.** Estimated peak time hours of five circadian genes for different genders and light conditions using the CircaCompare method.

|  | LD-F (hours) | LD-M (hours) | DD-F (hours) | DD-M (hours) |
| --- | --- | --- | --- | --- |
| *cyc* | 2.96 | 6.33 | na | 6.49 |
| *clk* | 9.87 | 10.52 | 4.29 | 5.40 |
| *tim* | 15.42 | 15.00 | 19.28 | 19.93 |
| *per* | 20.19 | 19.74 | 23.40 | 23.81 |
| *cry2* | 21.98 | 22.03 | na | 0.45 |

**Note:** LD-F or LD-M, heads sampled from female or male adults under 14 hours light: 10 hours dark (LD); DD-F or DD-M: heads sampled from female or male adults under the 3rd day of constant darkness (DD) after 7 days of adult entrainment to LD.

## Discussion

*S. frugiperda* is known to be active in tropical and subtropical regions (Kenis et al., 2022), with behaviors such as calling, mating, and ovipositing predominantly occurring after sunset (Hanniger et al., 2017; Miller et al., 2024; Pashley et al., 1992; Schofl et al., 2009). Nocturnal activity including eclosion behavior likely helps moths mitigate the adverse effects of high daytime temperatures and low humidity on physiological processes (He et al., 2021; Malekera et al., 2022). The eclosion of most insects occurs during specific time windows within a day, exhibiting circadian rhythmicity gated by the circadian clock (Mark et al., 2021; Wegener et al., 2024). Here, concentrated eclosion during the early dark phase also likely enhances survival by reducing exposure to diurnal avian predators (Pashley et al., 1992; Schofl et al., 2009). Besides, the circadian rhythm of adult eclosion (emergence) is a classic behavioral output that has been instrumental in defining the core properties of circadian clocks and identifying clock genes (Wegener et al., 2024). To date, the characteristics of eclosion behavior and the expression patterns of core clock genes in *S. frugiperda* have been poorly investigated (Lv et al., 2023). In this study, using a customized monitoring system, we further examined potential sex-dimorphic differences in both diel and circadian eclosion rhythms, as well as in the expression patterns of core clock genes. Our data provide compelling evidence for sexually dimorphic circadian organization at both behavioral and molecular levels in a migratory insect, suggesting that sex-dimorphic clock architecture may underlie divergent ecological strategies, including differences in eclosion behavior and migratory traits.

The light-dark cycle serves as a strong zeitgeber for diel eclosion rhythm (Merlin, 2009; Stanewsky, 2002). When entrained to a summer-like light-dark (LD) cycle, both female and male *S. frugiperda* exhibited a consistent eclosion peak shortly after lights-off at the light-dark transition, revealing a species-specific window for emergence timing (Bertossa et al., 2010; Nartey et al., 2020; Wang et al., 2023a). Here, although an anticipation of eclosion behavior to light-dark transition was observed before the peak shortly after light-off, the masking effect can’t be excluded when comparing the eclosion pattern with those under DD (Beer and Helfrich-Foerster, 2020; Bidell et al., 2024). The observed sex-specific differences in diel eclosion distribution may result from differences in clock gene mesor (i.e., the expression level) (Wei et al., 2025; Zhang et al., 2017), sexually dimorphic masking effects, or a combination of both, although the underlying mechanisms remain to be elucidated.

Across successive DD cycles, the eclosion peak shifts later each day, indicating an endogenous free-running period (τ) longer than 24 hours, a characteristic of circadian rhythms in the absence of external zeitgebers (Numata and Tomioka, 2023). Our recent work revealed a τ for circadian eclosion of *S. frugiperda* exceeding 24 hours with mixed females and males (Lv et al., 2023). In this study, we closely examined the potential sexually dimorphic circadian eclosion behavior and observed a consistent τ over 24 hours for both females and males, with a significant shift in timing between the sexes. Since knockdown of per has been shown to cause a phase shift in the circadian rhythm of courtship vibrations in *Nilaparvata lugens* (Wei et al., 2025), the observed difference in emergence timing between sexes may be attributed to sex-specific differences in per transcript mesor. Furthermore, the significant phase shift between diel and circadian eclosion profiles may be explained by corresponding phase differences in tim and per expression between LD and DD conditions in both females and males. Similar correlations between eclosion phenotypes and phase shifts in clock gene expression have been reported in *Danaus plexippus* (Zhang et al., 2017) and *Delia antiqua* (Miyazaki et al., 2016).

At the molecular level, we observed sex-dimorphic rhythmic parameters for the core clock genes *cyc*, *clk*, *tim*, *per*, and *cry2* in *S. frugiperda*, indicating sexual dimorphism in the regulation of behavioral rhythms by circadian clockwork. Under LD conditions, the expression patterns of *tim*, *per*, and *cry2* were consistent with those reported by Hänniger et al for *S. frugiperda* (Hanniger et al., 2017), as well as with expression profiles observed in *Agrotis ipsilon* and *Danaus plexippus* (Li et al., 2025). This similarity suggests that the expression patterns of *tim*, *per*, and *cry2* in the heads of migratory insects may be evolutionarily conserved. However, in contrast to our findings, Hänniger *et al* reported that *cyc* and *clk* lacked rhythmicity in the rice-and corn-strain *S. frugiperda* under LD condition (Hanniger et al., 2017), which may be attributed to hybridization between strains during the invasion of China (Wang et al., 2024). CircaCompare analysis revealed that under both LD and DD conditions, the mesor of *cyc*, *clk*, and *per* in males was significantly higher than in females, with *tim* and *cry2* showing significantly higher mesor in males only under LD conditions. Hänniger et al noted that *cyc*, *clk*, and *per* are located on the Z chromosome in *S. frugiperda*, which follows a ZW sex-determination system (females: ZW; males: ZZ) (Hanniger et al., 2017). The presence of two Z chromosomes in males may contribute to a gene dosage effect (Dayton and Dopman, 2024; Goh, 2022), resulting in significantly higher mesor for *cyc*, *clk*, and *per* in males compared to females under both conditions. However, the mesor in males was not exactly double that in females, suggesting potential compensatory regulatory mechanisms. As transcriptional activators, the highly expressed *clk* and *cyc* genes may enhance the expression of downstream genes *tim* and *cry2* (Brady et al., 2021), thereby contributing to the overall higher transcriptional levels of core clock genes in males under LD conditions.

Sexual dimorphism in circadian systems has emerged as a critical aspect of chronobiology, with growing evidence indicating that males and females often differ in their circadian behaviors, clock gene expression, and physiological rhythms (Bailey and Silver, 2014; Duffy et al., 2011; Force et al., 2024; Helfrich-Forster, 2000; Joye and Evans, 2022; Krizo and Mintz, 2015; Rund et al., 2012; van Doorn et al., 2024; Walton et al., 2022). Our findings provide empirical evidence supporting the hypothesis that sexually dimorphic circadian regulation can facilitate allochronic differentiation within populations, as proposed in recent theoretical models (van Doorn et al., 2024). Here, significant sex-specific differences in the mesor and phase of core clock gene expression, accompanied by distinct diel eclosion profiles between males and females suggest that circadian architecture differs between sexes, potentially generating sex-specific temporal windows for critical life history processes such as mating or migration. This is consistent with simulation studies indicating that sex-specific expression of circadian genes can mitigate the homogenizing effects of sexual selection—particularly male-mating competition—thereby maintaining or even promoting chronotype divergence in sympatry (van Doorn et al., 2024). Given the species’ rapid global invasion and evidence of strain hybridization (Kenis et al., 2022; D. Wang et al., 2023b; L. Zhang et al., 2023), the emergence of sexually dimorphic circadian traits in invasive populations may reflect an adaptive response that enables temporal niche partitioning. Future research expanding to geographically distinct populations—especially between native and invasive ranges—will be essential to determine whether such sex-specific circadian traits are conserved or subject to local ecological pressures. Moreover, integrating additional circadian-regulated behaviors—such as mating (Zhang et al., 2022) and flight rhythms (He et al., 2021), under variable photoperiods and simulated natural conditions will be crucial to understanding how sex-and population-level circadian divergence contributes to ecological adaptation and the early stages of allochronic speciation.

## Acknowledgments

This research is funded by the National Natural Science Foundation of China (grant nos. 32172414 and 32472547), the Natural Science Foundation of Jiangsu Province (BK20221510).

## Supplementary Information

**Fig. S1.**
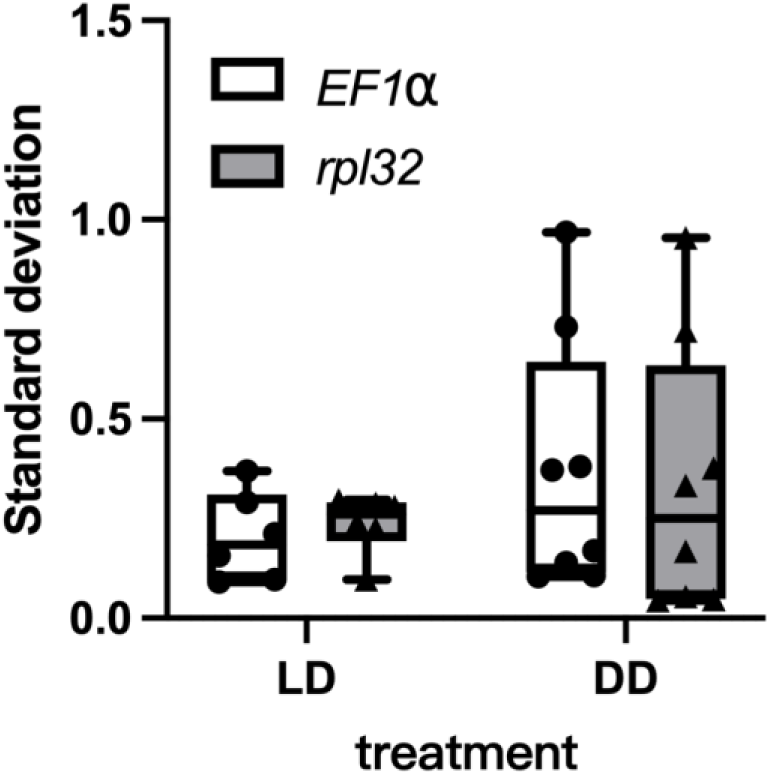
Standard deviation (SD) of two reference genes *EF1⍺* and *rpl32* in all 84 samples under female versus male.

## Notes

### Competing Interest Statement

The authors have declared no competing interest.

